## Supplementary figures and images for "Three new integration vectors and fluorescent proteins for use in the opportunistic human pathogen *Streptococcus pneumoniae*"

### Supplementary Figure 1

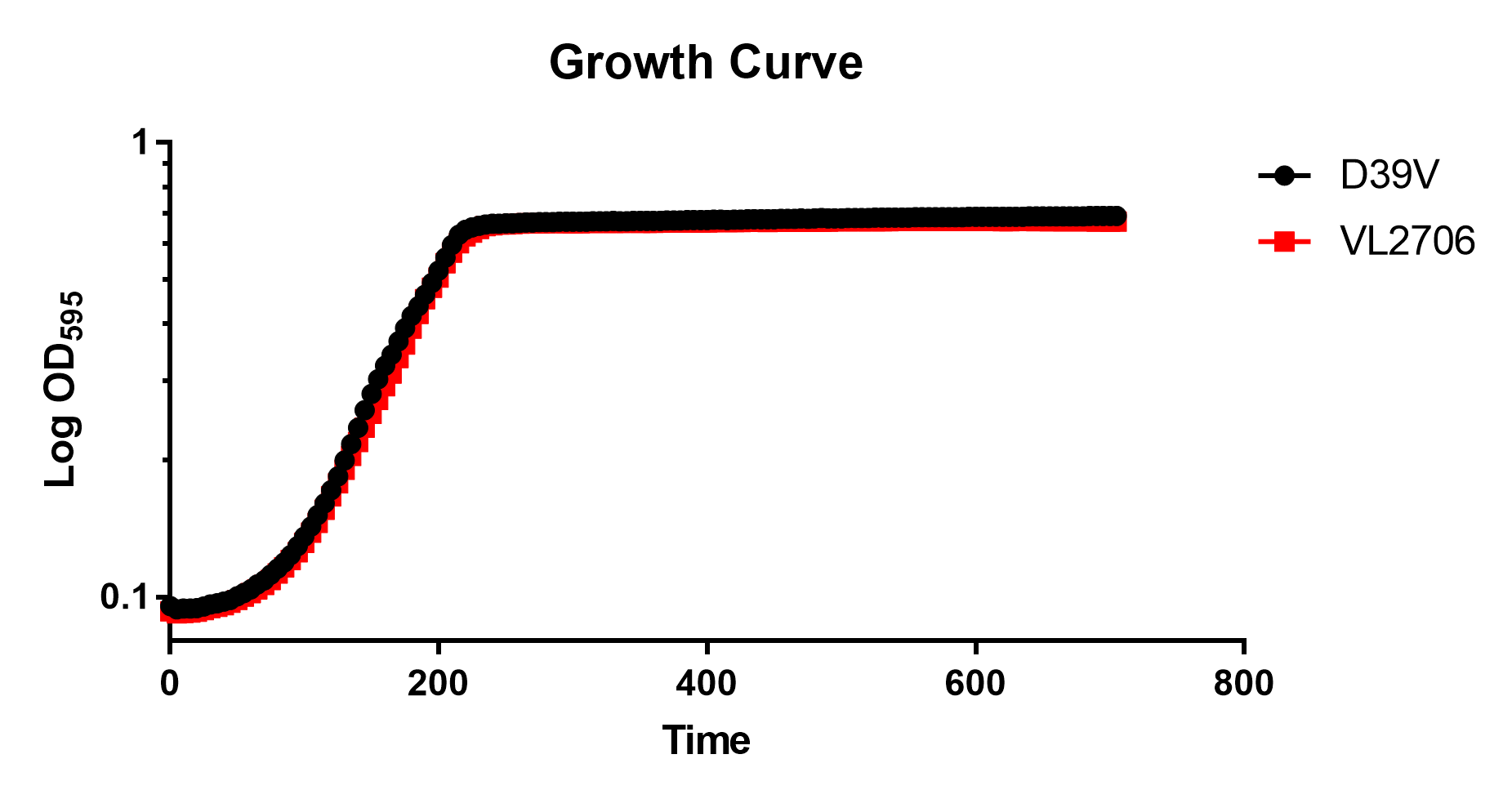

### Supplementary Figure 2

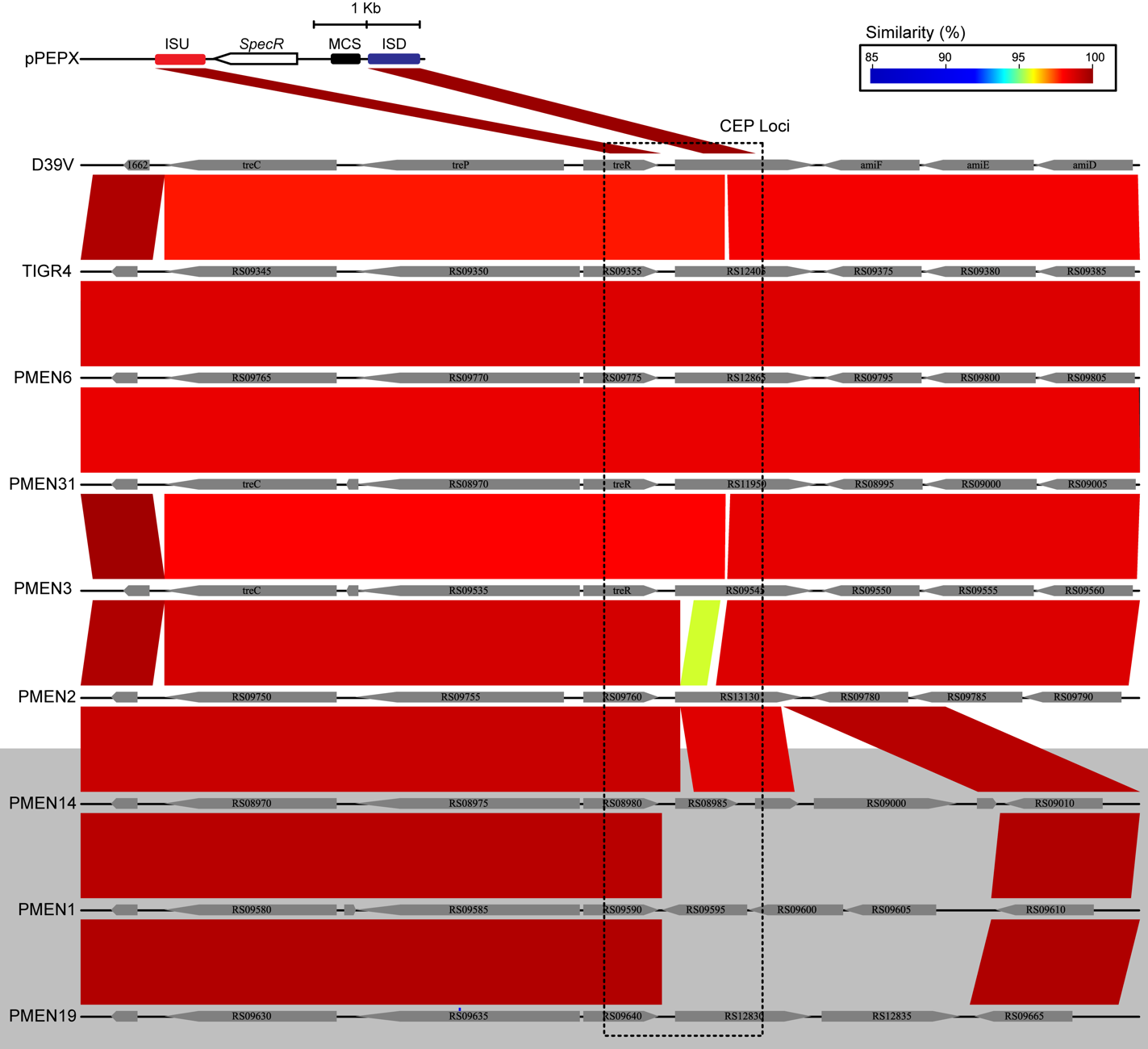

### Supplementary Figure 3

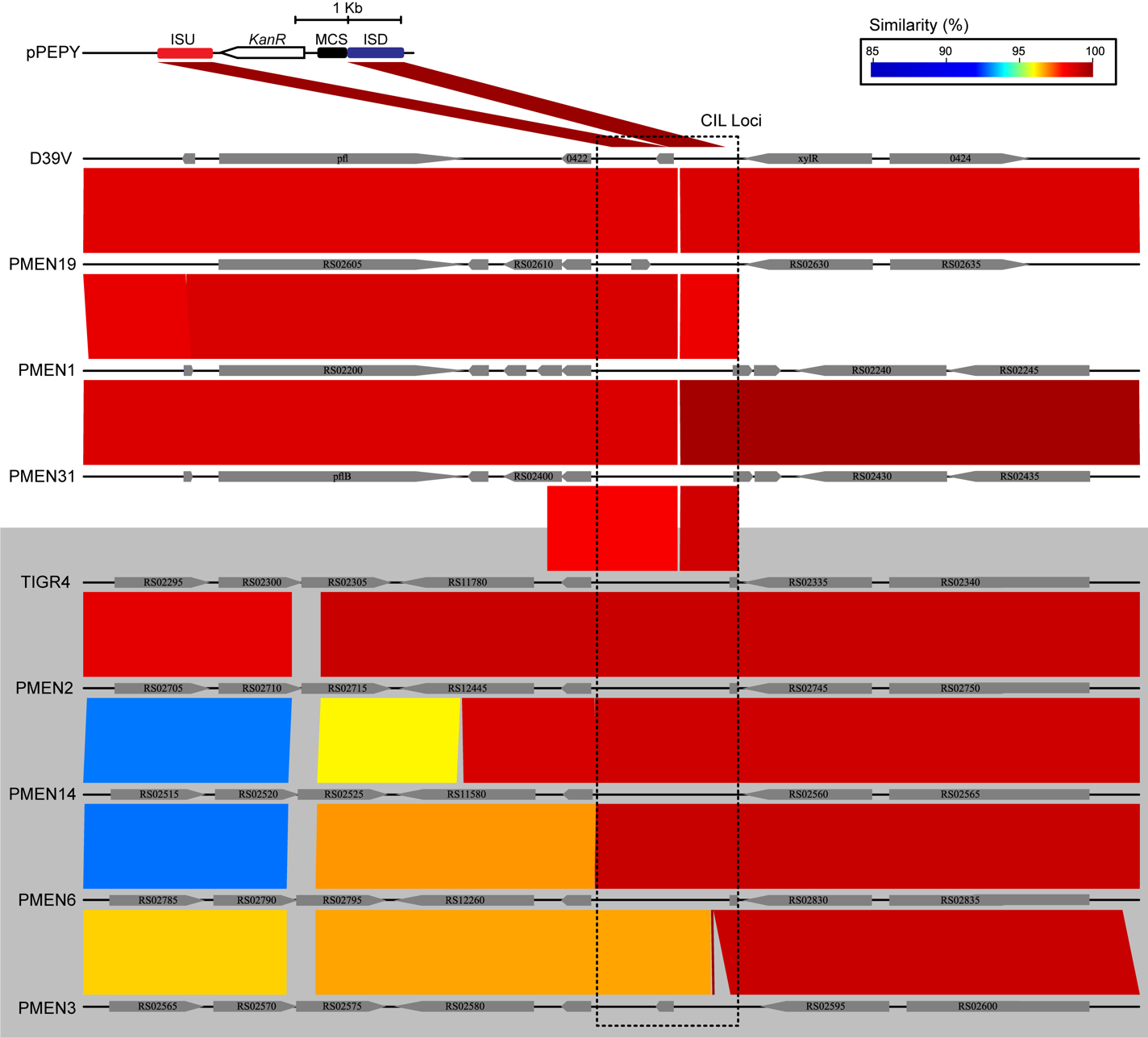

### Supplementary Figure 4

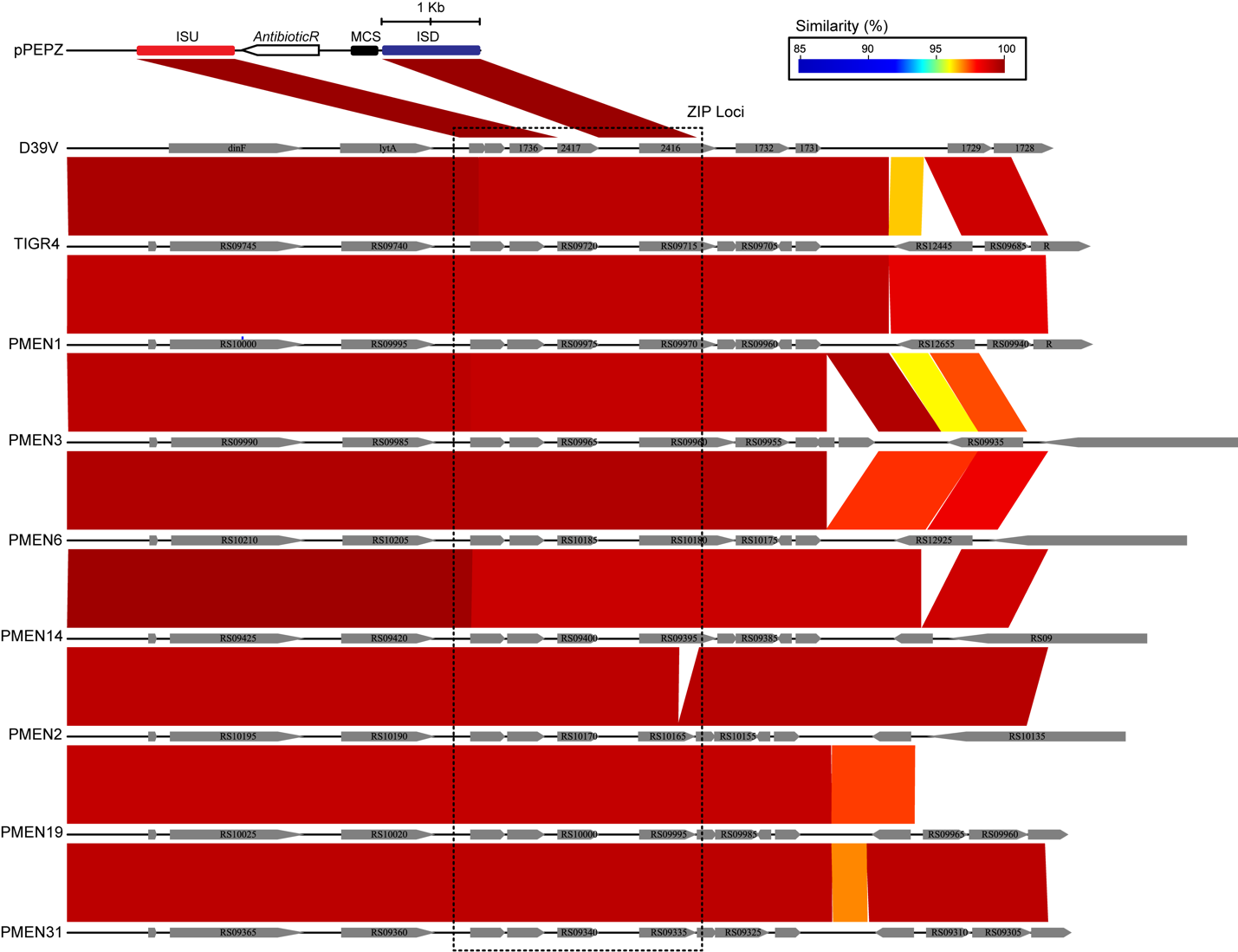
