## Supplementary Figure Legends for "Three new integration vectors and fluorescent proteins for use in the opportunistic human pathogen *Streptococcus pneumoniae*"

Figure S1: Growth curve of parent strain D39V and VL2706. Strain VL2706 has pPEPX, pPEPY, and pPEPZ chromosomally integrated each expressing an extra copy of a fluorescently fused protein that has a distinct localization. As seen in the growth curve the growth rate and maximal bacterial density is not different from the parent strain D39V.

Figure S2: Comparative alignment for CEP (pPEPX) locus and its flanking regions from various *S. pneumoniae* strains. Homology region for CEP locus for pPEPX integration from complete genomic sequences of PMEN strains were aligned and compared based on bl2seq (BLASTn). Color code represents nucleotide sequence similarity. The five strains (TIGR4, PMEN6, PMEN31, PMEN3 and PMEN2) harbor conserved CIL homologous loci (highlighted by dashed frame) as well as D39V. Meanwhile, three strains (PMEN14, PMEN1 and PMEN19) have less sequence similarity, which might results in less transformability. Reference genomic sequences of each PMEN strains were obtained from NCBI. Strain names and accession number for each PMENs are following: D39V (GCA_003003495.1); TIGR4 (TIGR4; ATCC BAA-334, GCA_000006885.1); PMEN1 (ATCC 700669, GCA_000026665.1); PMEN2 (670-6B, GCA_000147095.1); PMEN3 (4041STDY6836167, GCA_900476445.1); PMEN6 (Hungary19A-6, GCA_000019265.1); PMEN14 (Taiwan19F-14, GCA_000019025.1); PMEN19 (70585, GCA_000018965.1); PMEN31 (OXC141, GCA_000210955.1 ).

Figure S3: Comparative alignment for CIL (pPEPY) locus and its flanking regions from various *S. pneumoniae* strains. Homology region for CIL locus for pPEPY integration from complete genomic sequences of PMEN strains were aligned and compared based on bl2seq (BLASTn). Color code represents nucleotide sequence similarity. The three strains (PMEN1, PMEN19 and PMEN31) harbor conserved CIL homologous loci (highlighted by dashed frame) as well as D39V. Meanwhile, five strains (TIGR4, PMEN2, PMEN14, PMEN6 and PMEN3) have less sequence similarity, which might results in less transformability. Reference genomic sequences of each PMEN strains were obtained from NCBI. Strain names and accession number for each PMENs are described in Fig S2.

Figure S4: Comparative alignment for ZIP (pPEPZ) locus and its flanking regions from various *S. pneumoniae* strains. Homology region for ZIP locus for pPEPZ integration from complete genomic sequences of PMEN strains were aligned and compared based on bl2seq (BLASTn). Color code represents nucleotide sequence similarity. All of the tested nine PMEN strains harbor highly conserved ZIP homologous loci (highlighted by dashed frame) as well as D39V. Reference genomic sequences of each PMEN strains were obtained from NCBI. Strain names and accession number for each PMENs are described in Fig S2.

Table S1: List of oligos used for strain construction. Italics indicates overlapping region and bolded indicates restriction sites.

Table S2: List of *Streptococcus pneumoniae* strains tested homology to allow for chromosomal integration of the PEP vectors.
